# Distinct tissue-dependent composition and gene expression of human fetal innate lymphoid cells

**DOI:** 10.1101/2024.03.12.583977

**Authors:** Inga E. Rødahl, Martin A. Ivarsson, Liyen Loh, Jeff E. Mold, Magnus Westgren, Danielle Friberg, Jenny Mjösberg, Niklas K. Björkström, Nicole Marquardt, Douglas F. Nixon, Jakob Michaëlsson

## Abstract

The human fetal immune system starts to develop in the first trimester and likely plays a crucial role in fetal development and maternal-fetal tolerance. Innate lymphoid cells (ILCs) are the earliest lymphoid cells to arise in the human fetus. ILCs consists of natural killer (NK) cells, ILC1s, ILC2s, and ILC3s that all share a common lymphoid origin. Here, we studied fetal ILC subsets, mainly NK cells and ILC3s and their potential progenitors, across human fetal tissues from the first and second trimesters. Our results show that fetal ILC subsets have distinct distribution, developmental kinetics, and gene expression profiles across human fetal tissues. Furthermore, we identify a putative CD34^+^RORγt^+^Eomes^+/−^ ILC progenitor population exclusively present in fetal intestine, indicating that tissue-restricted development of ILCs could contribute to the variation in ILC composition and gene expression between tissues.

## Introduction

Natural killer (NK) cells and other innate lymphoid cells (ILCs) are rapidly responding immune cells found in lymphoid and non-lymphoid tissues (1). Human ILCs are divided into three major groups: group 1 ILCs consisting of cytotoxic and IFN-γ producing Eomes^+^Tbet^+/−^ NK cells (2) and IFN-γ-producing Eomes^−^Tbet^+^ ILC1 (3), group 2 ILCs consisting of IL-5 and IL-13 producing GATA-3^+^ ILC2 cells (4, 5), and group 3 ILCs consisting of IL-17 and IL-22 producing RORγt^+^ cells (6-9). Group 3 ILCs also include lymphoid tissue inducer (LTi) cells (10, 11), shown to be important for lymphoid tissue formation in mice. In humans, ILC3s with LTi activity are enriched in CD304^+^ (neuropilin, *NRP1*) ILC3s (12). ILCs are also commonly identified by expression of combinations of cell surface proteins. Most studies define NK cells as Lin^−^CD56^+^CD127^−^ cells and non-NK cell ILCs as Lin^−^CD127^+^ cells. Among non-NK cell ILCs, ILC3 are defined as Lin^−^CD127^+^CD117^+^NKp44^+/−^ cells, ILC2 as Lin^−^CD127^+^CRTH2^+^ cells, and ILC1 as Lin^−^CD127^+^CD117^−^NKp44^+/−^ cells. Variation in the definitions of ILC subsets between studies, including use of different lineage markers, has to some extent made it hard to compare results across studies. For example, CD7 (13, 14) and CD11b (15) expression have been respectively used as an inclusion and exclusion criteria in some studies, whereas they are not used in other studies.

NK cells, ILC2s, and ILC3s can be detected in fetal tissues already during the first trimester (6, 15-17), whereas ILC1s appear to develop later, possibly after birth (3). NK cells have been detected in fetal liver and skin as early as post conceptional week (PCW) 6 (16-18), and functional CD16^−^ and CD16^+^ NK cells capable of responding to target cells and cytokines are present in second trimester fetal tissues (19, 20). In addition to NK cells, ILC3s are abundant in fetal tissues (13, 15, 17, 18), with considerable heterogeneity in terms of both protein and RNA expression (13, 15, 18). CD304^+^ ILC3s are abundant in fetal spleen and lymph nodes and are of particular interest given their suggested role as LTi cells (12).

Each of the ILC groups have different distribution in adult tissues. Generally, NK cells predominate among ILCs in most adult tissues studied, including liver, lung, uterus, spleen, and blood (21-23), although ILC3s are enriched in the intraepithelial compartment of the ileum (24). In addition, the different ILC subsets have distinct gene expression profiles in different adult tissues (24, 25), potentially reflecting subset-specific migratory patterns and/or tissue-specific imprinting.

A number of studies have explored the stages at which different groups of ILCs diverge from each other (14, 15, 26-29). A fraction of the common innate lymphoid progenitor (CILP) cells, defined as CD34^+^CD117^+^IL1R1^+^RORγt^+^, in tonsils can generate all groups of ILCs, including NK cells (28). Similarly, a fraction of CD34^−^CD117^+^CD127^+^RORγt^−^ cells (ILCPs) in cord blood, adult blood, and lung can differentiate into all groups of ILCs (14), indicating that the divergence of ILC subsets may occur late in maturation. Downstream of CILPs and ILCPs, CD34^−^CD117^+^CD56^+^ cells in tonsil can give rise to both NK cells and ILC3s, but not ILC2s (30). However, a CD34^+^ NK cell-restricted progenitor in cord blood, defined as CD34^+^CD45RA^+^CD7^+^CD10^+^CD127^−^ cells, has also been identified, suggesting that at least some NK cells might diverge from the common ILC development at an earlier stage (27). There is considerable heterogeneity within each of the described progenitor populations, leaving room for more refined studies of ILC development. In addition, it remains possible that ILCs develop differently depending on tissue location *in vivo*, and that even phenotypically similar ILCs can develop from distinct, possibly tissue-restricted, progenitors.

Although it is known that ILCs develop early in the human fetus, there is a lack of studies examining ILCs at subset level throughout fetal development, spanning gestation and across multiple tissues, using combined analysis of key transcription factors and cell surface markers. Moreover, little is known regarding gene and protein expression differences among ILC subsets across fetal tissues. In order to address these questions, we analyzed ILC subsets and progenitors from human fetal tissues using flow cytometry and RNA sequencing. We demonstrate distinct ILC subset distribution and gene expression profiles of human ILCs across fetal tissues. Collectively, our findings enhance our understanding of ILC ontogeny, composition and dynamics in various fetal tissues, and shed light on novel putative tissue-restricted ILC progenitor populations.

## Results

### Fetal ILCs have distinct subset composition and developmental kinetic dependent on tissue location

We first analyzed human fetal ILCs in liver, lung, intestine, bone marrow and skin during the first and second trimester of gestation. Using multicolor flow cytometry, we analyzed expression of transcription factors important for ILC development and function (1), including Eomes, T-bet, RORγt, GATA-3, and PLZF, together with expression of cell surface receptors. We identified NK cells as Lin^−^ (CD3^−^CD14^−^CD19^−^)CD34^−^Eomes^+^ cells, group 3 ILCs as Lin^−^CD34^−^Eomes^−^RORγt^+^ cells, and group 2 ILCs as Lin^−^CD34^−^Eomes^−^RORγt^−^GATA-3^+^ cells (**Fig. 1A, Fig. S1A**) in first trimester fetal tissues. Consistent with previous studies (3), virtually no *bona fide* ILC1 cells defined as Lin^−^CD34^−^Tbet^+^Eomes^−^CD127^+^ could be identified, while Tbet expression was readily detected in Eomes^+^ NK cells (**Fig. S1B**). In contrast to mice (31), and consistent with studies of adult ILCs (30, 32), all fetal ILC subsets expressed PLZF (**Fig. 1B**) in all tissues analyzed (**Fig. 1C**), albeit at higher levels in group 2 and 3 ILCs compared to NK cells (**Fig. 1B-C**).

**Figure 1.**
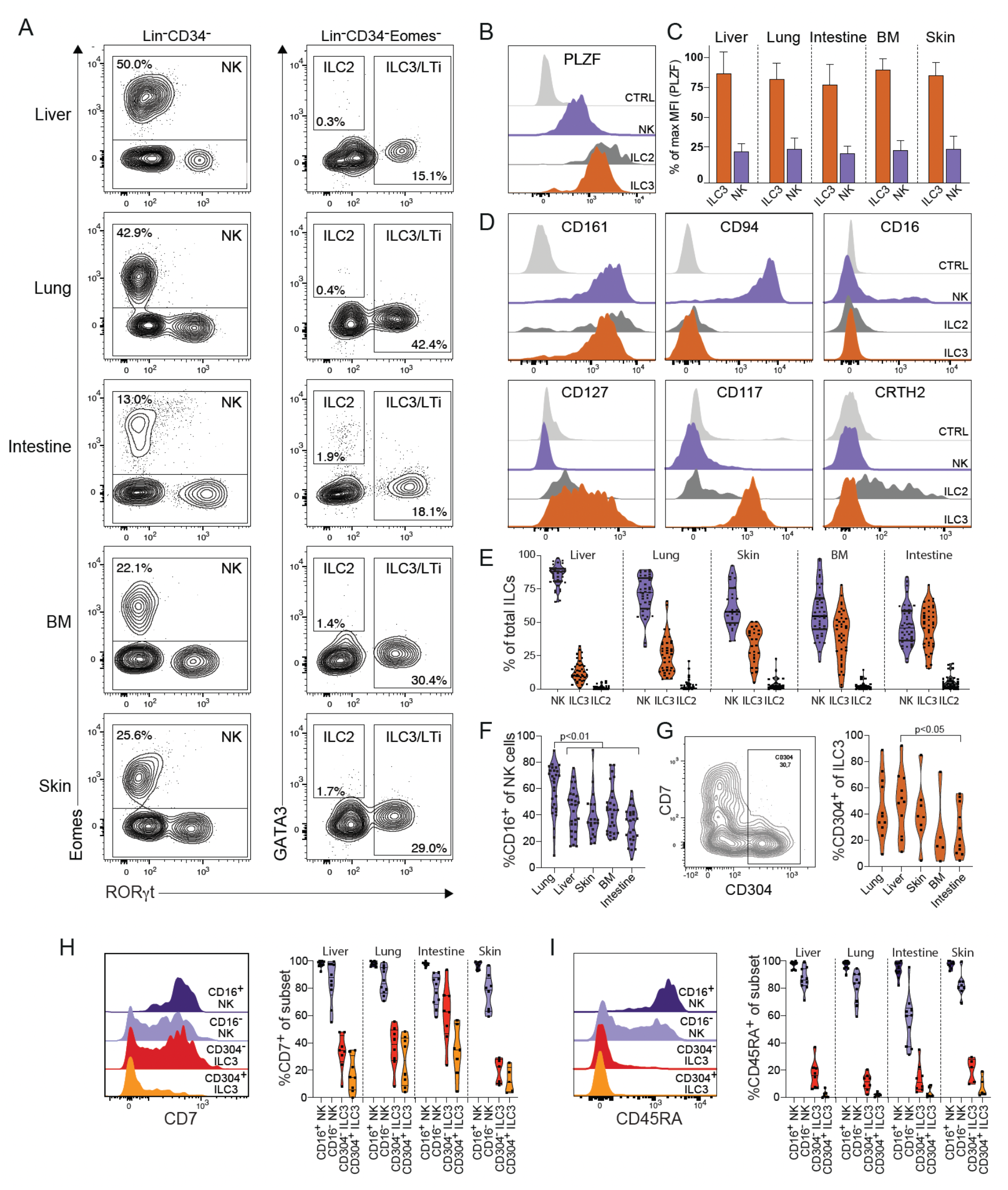
Characterization of fetal ILC subsets across tissues using flow cytometry. (**A**) Representative staining and gating of human NK cells (Lin^−^CD34^−^Eomes^+^), ILC2 (Lin^−^CD34^−^RORγt^−^GATA3^+^) and ILC3 (Lin^−^CD34^−^Eomes^−^RORγt^+^) in fetal liver, lung, intestine, bone marrow, and skin at PCW 9.5. (**B**) Representative histogram showing expression of PLZF in intestinal NK cells, ILC2s, and ILC3s at PCW 9.5. ILC subsets defined as in (A). (**C**) Relative fluorescence intensity in ILC3s and NK cells across tissues. Bars represent mean with SD. *n* = 14 (bone marrow (BM)), *n* = 11-13 (intestine, liver, and lung), *n* = 7-8 (skin) at PCW 8-11. (**D**) Representative flow cytometry histogram showing expression of CD161, CD94, CD16, CD127, CD117 and CRTH2 on NK cells, ILC2s, and ILC3s in fetal intestine at PCW 9.5. (**E**) Violin plots showing frequency of ILC subsets of total ILCs across fetal liver, lung, skin, bone marrow, and intestine. Bars indicate mean, *n* = 38-40, *n* = 27 (skin only). (**F**) Violin plots showing frequency of CD16^+^ NK cells among total NK cells across tissues. Statistical analysis using one-way related measures ANOVA, n = 39-40 (liver and lung), n = 35-36 (intestine, BM), n = 26 (skin). (**G**) Representative staining of CD304 and CD7 on ILC3s (left, defined as in A) at PCW 10 and frequency of CD304^+^ ILC3 of total ILC3s across tissues (right). Statistical analysis using one-way related measures ANOVA, paired tissues *n* = 11 (liver, lung, and gut), *n* = 8 (skin), *n* = 5 (BM). (**H-I**) Representative histograms and frequencies of (**H**) CD7^+^ cells and (**I**) CD45RA^+^ cells, among CD16^+^ and CD16^−^ NK cells and CD304^−^ and CD304^+^ ILC3s **(*n* = 5-10)**.

The ILC subsets identified by analyses of transcription factor expression had a cell surface phenotype consistent with previous reports (1). Virtually all ILCs expressed CD161, only Eomes^+^ NK cells expressed CD94, group 3 ILCs expressed CD117, group 2 and 3 ILCs expressed CD127, and a majority of group 2 ILCs expressed CRTH2 (**Fig. 1D**). However, in contrast to adult NK cells (33), only a subset of CD16^−^ and CD16^+^ NK cells expressed NKp80 (**Fig. S1C**).

NK cells and group 3 ILCs were the dominant ILC subpopulations across all tissues, whereas group 2 ILCs constituted less than 10% of the total ILCs (**Fig. 1E**). NK cells made up the majority of ILC in liver, lung, skin, and bone marrow, whereas NK cells and ILC3 were equally frequent in the intestine (**Fig. 1E**). The frequency of NK cells among ILCs increased significantly over gestation in liver and lung at the expense of a decrease in ILC3s, whereas no significant changes could be detected over time in the intestine (**Fig. S1E**). Confirming previous studies (16, 17, 34), ILCs developed several weeks before T cells could be detected in peripheral tissues (**Fig. S1F**). Given the variation in NK cell and ILC3 frequencies between fetal tissues, we next determined whether the composition of each ILC subset also varied. NK cells are commonly divided into CD56^bright^CD16^−^ and CD56^dim^CD16^+^ NK cells, with distinct functions and tissue distribution (35). Similarly, CD304^+^ and CD304^−^ ILC3s have distinct tissue distribution and gene expression profiles (12). The frequency of CD16^+^ NK cells was highest in lung (**Fig. 1F**), and the frequency of CD304^+^ ILC3s was highest in liver (**Fig. 1G**), while the frequencies of both subsets were lowest in intestine (**Fig. 1F-G**).

Expression of CD7 and CD45RA have been reported to be downregulated both in highly differentiated adaptive NK cells (32) and in differentiated ILC3 (36). Virtually all CD16^+^ NK cells, and the majority of CD16^−^ NK cells, expressed CD7 and CD45RA in all fetal tissues (**Fig. 1H-I**). In contrast, only a small fraction of CD304^+^ and CD304^−^ ILC3s expressed CD7 or CD45RA, except for CD304^−^ ILC3s in the intestine (**Fig. 1H-I**). Notably, the expression of CD7 on ILC3s increased with gestation in liver, lung, and intestine (**Fig. S1D**). Taken together, NK cells and ILC3s make up the vast majority of ILCs during the first and second trimester but vary in composition between tissues and over gestational age.

### Fetal ILC subsets have tissue-specific transcriptional profiles

To increase the depth of our analysis of fetal ILCs, we analyzed tissue-specific gene expression patterns within ILC subsets. To this end, we analyzed RNA expression in FACS-sorted CD16^+^ and CD16^−^ NK cells, as well as CD304^+^ ILC3s, from human fetal liver, intestine, skin, and lung (gating strategy in **Fig. S2A**). As expected, subset identity contributed to the largest variation in gene expression in principal components analysis (PCA) (**Fig. S2B**).

Pairwise differential gene expression analysis between tissues for each subset revealed 218 differentially expressed genes (padj < 0.01, log2foldchange > 1) in CD304^+^ ILC3s, and 164 and 128 differentially expressed genes in CD16^−^ and CD16^+^ NK cells, respectively (**Fig. S3A-B, Supplementary data 1**). We next focused on the unique gene expression profile of each tissue and ILC subset (**Fig. 2A-C**). NK cells in skin and liver had more distinct transcriptional profiles compared to NK cells in fetal intestine and lung (**Fig. 2A-B, S3**). CD304^+^ ILCs had distinct transcriptional profiles across all four tissues (**Fig. 2C**). The tissue-specific gene expression profiles were largely unique for each subset (**Fig. 2D**), indicating that local factors within each distinct tissue does not drive a conserved gene expression pattern in all ILC subsets.

**Figure 2.**
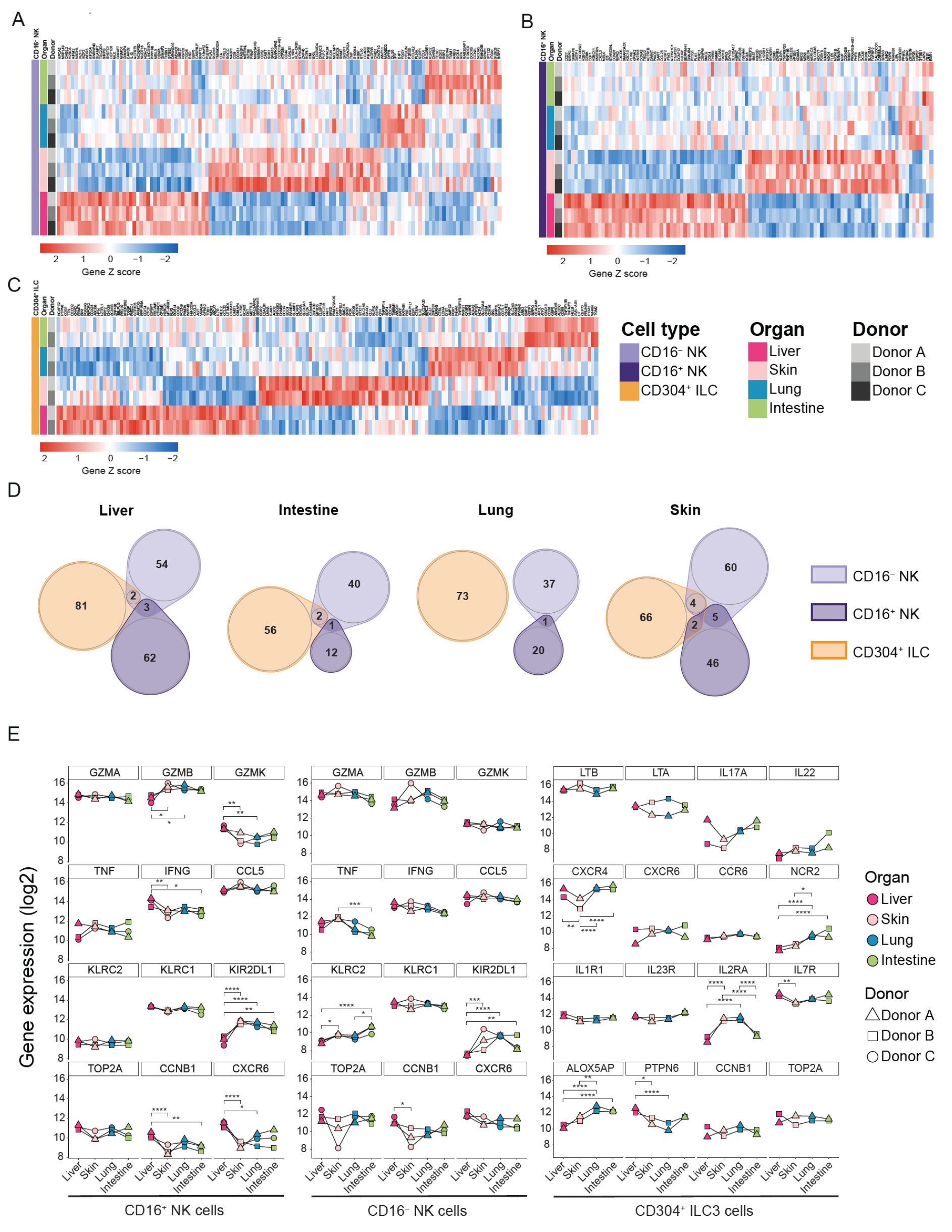
Fetal ILC subsets have tissue-specific transcriptomic profiles. (**A-C**) Heatmaps of unique differentially expressed genes from pairwise comparison of fetal tissues. Genes are grouped by tissue and by expression level, visualized by z-score. Only genes uniquely differentially expressed by one tissue are included. (**A**) CD16^−^ NK cells (*n* = 3), (**B**) CD16^+^ NK (*n* = 3) cells, and (**C)** CD304^+^ ILCs (*n* = 2). Differentially expressed genes from pairwise comparisons with padj < 0.01 and log2 fold change > 1 are shown. (**D**) Euler diagram depicting overlap of differentially expressed genes from each subset in the same tissue. (**E**) Normalized gene expression of genes of interest in CD16^+^ NK cells, CD16^−^ NK cells, and CD304^+^ ILC3s. \**padj* < 0.05, \*\**padj* < 0.01, \*\*\**padj* < 0.001, \*\*\*\**padj* < 0.0001 from pairwise comparisons using DESeq2. Matched tissues from 2-3 donors.

Focusing on gene expression unique for each tissue could potentially mask genes expressed at higher levels in multiple tissues by a given subset (**Fig. 2A-C**). We therefore explored the overall gene expression and focused on canonical NK cell and ILC3 genes (**Fig. S3**, supplementary data 1). Analysis of NK cells revealed several differentially expressed genes relevant for NK cell regulation and function, including activating and inhibitory receptors (*KIR*s and *KLRC2*) and effector molecules (*GZMB, IFNG, TNF*). Notably CD16^+^ NK cells in fetal liver expressed lower levels of *GZMB* and *KIR2DL1* coupled with higher expression of *GZMK, IFNG*, and *CXCR6* compared to other tissues (**Fig. 2E**). In contrast, CD16^−^ NK cells expressed similar levels of *GZMB, GZMK* and *IFNG* across tissues, whereas they expressed less *KLRC2* and *KIR2DL1* in liver. Lastly, CD16^+^ NK in liver expressed higher levels of *CCNB1* (Cyclin B1) compared to other tissues, indicative of more proliferation in liver.

CD304^+^ ILC3s expressed high levels of *LTB* and *LTA* in all tissues consistent with their proposed function as LTi-like cells. The canonical ILC3 cytokine receptors *IL1R1* and *IL23R* were also expressed at similar levels across all tissues. In contrast, expression of several genes regulating ILC3 function or migration, including *NCR2, IL2RA, IL7R, PTPN6, ALOX5AP*, and *CXCR4*, varied significantly between tissues. For example, *NCR2* (NKp44) associated with ILC3 maturation (36, 37), and *ALOX5AP, ZNF331, RGS1* and *RGS2*, associated with tissue residency (21) were significantly upregulated in lung and intestine relative to liver (**Fig. 2E, S3A**). While there were no statistically significant differences in expression of *IL17A* and *IL22*, there was a trend of higher expression of these cytokines in the intestine (**Fig. 2E**). Together, our analysis suggested that human fetal ILC subsets have transcriptomic profiles unique to both tissue and subset.

### Human fetal intestine contains putative CD34^+^RORγt^+^Eomes^+/−^ ILC progenitors

To explore whether the tissue-specific gene expression patterns could be due to development of ILCs from tissue-restricted ILC progenitors, we investigated putative CD34^+^ ILC-progenitors in fetal tissues. The frequencies of Lin^−^CD45^+^CD34^+^ cells were highest in fetal liver and intestine, in particular early in gestation, with substantially lower frequencies in bone marrow and lung (**Fig. 3A-B**).

**Figure 3.**
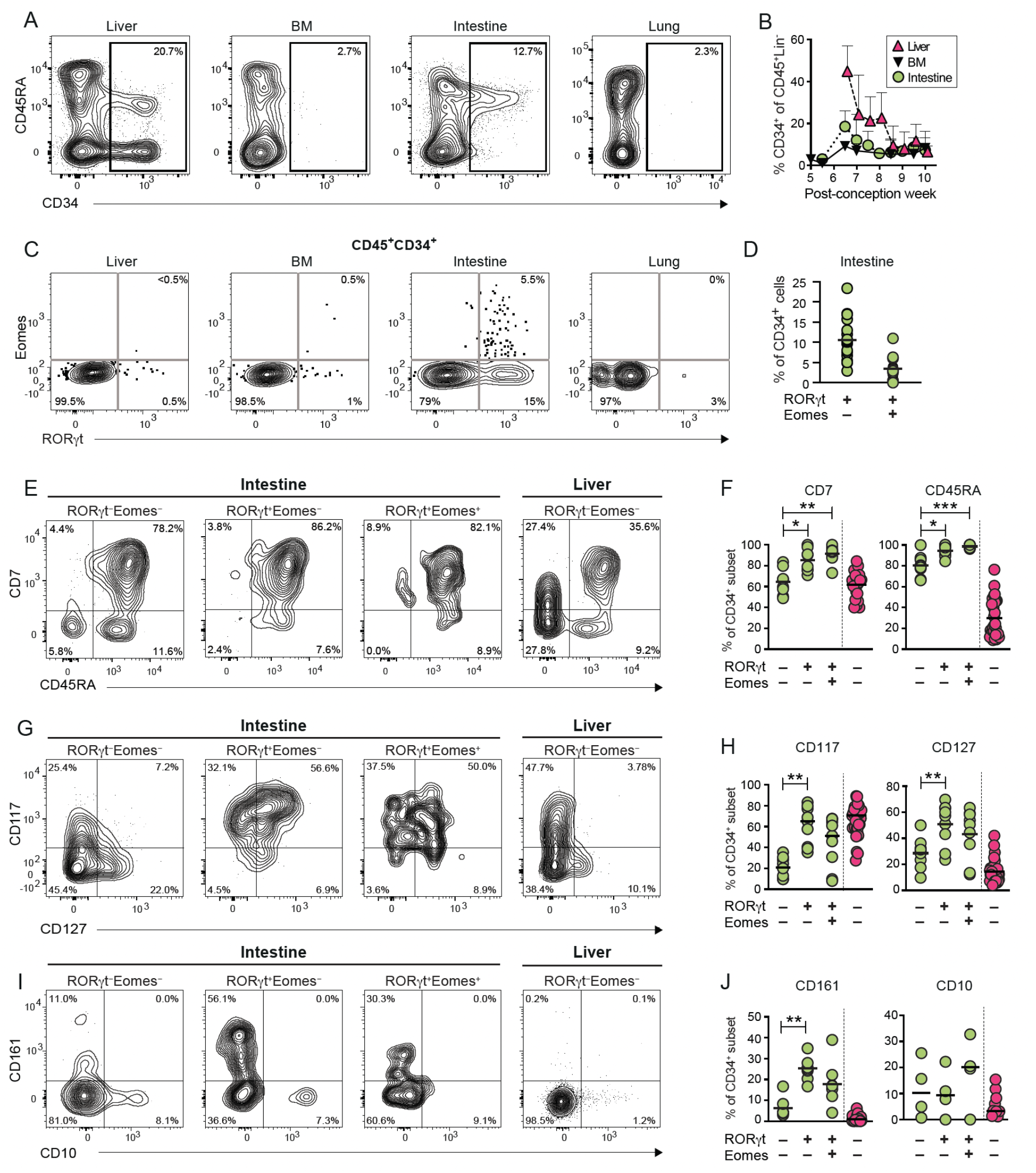
Fetal CD34^+^ progenitors co-express Eomes and RORγt in the intestine. (**A**) Representative contour plots of CD45RA and CD34 expression by Lin^-^CD45^+^ cells in fetal liver, bone marrow, intestine, and lung at PCW 7.5. (**B**) Mean frequency of CD34^+^ cells of Lin^-^CD45^+^ cells across gestational age in fetal liver (n = 24), bone marrow (n=13) and intestine (n=24). (**C**) Representative contour plots of Eomes and RORγt expression by Lin^-^ CD45^+^CD34^+^ cells in liver, bone marrow, lung, and intestine at PCW 9.5. (**D**) Frequency of RORγt^+^Eomes^−^ and RORγt^+^Eomes^+^ of CD34^+^ cells in intestine (n = 17). Bar shows mean frequency. (**E-J**) Representative contour plots (PCW 9.5) and summary of data for frequency of expression of (**E-F**) CD7 and CD45RA, (**G-H**) CD117 and CD127, and (**I-J**) CD161 and CD10 on CD34^+^RORγt^−^Eomes^−^, CD34^+^RORγt^+^Eomes^−^ and CD34^+^RORγt^+^Eomes^+^ cells in fetal intestine (green, n = 8-9), and on CD34^+^RORγt^−^Eomes^−^ cells in fetal liver (pink, n = 21-33), respectively. Statistical comparison by non-parametric ANOVA. \**padj* < 0.05, \*\**padj* < 0.01, \*\*\**padj* < 0.001.

CD34^+^RORγt^+^ progenitors have previously been described as ILC restricted in adult tonsil (28, 29), but it remains unknown if cells with a similar phenotype exist in fetal tissues. We identified CD34^+^RORγt^+^Eomes^−^ cells and CD34^+^RORγt^+^Eomes^+^ cells in the fetal intestine (**Fig. 3C-D**), whereas they were scarce or absent in first trimester fetal liver, lung, and bone marrow (**Fig. 3C**). Virtually all CD34^+^RORγt^+^Eomes^−^ and CD34^+^RORγt^+^Eomes^+^ cells expressed CD45RA and a large majority expressed CD7 (**Fig. 3E-F)**. Moreover, CD34^+^RORγt^+^Eomes^−^ cells more frequently expressed CD117, CD127 and CD161 compared to CD34^+^RORγt^−^Eomes^−^ cells, and a similar trend was observed for CD34^+^RORγt^+^Eomes^+^ cells (**Fig. 3G-J**). Finally, CD10, reported to define NK cell-restricted progenitors (27), was not more frequently expressed on CD34^+^RORγt^+^Eomes^−^ and CD34^+^RORγt^+^Eomes^+^ cells compared to CD34^+^RORγt^−^Eomes^−^ cells in the intestine (**Fig. 3I-J**). In summary, we identified potential pluripotent Lin^−^CD34^+^RORγt^+^Eomes^+/−^ ILC progenitor cells restricted to fetal intestine, indicative of tissue-specific ILC development.

### Eomes and RORγt are co-expressed in CD34^−^CD16^−^ fetal NK cells

CD34^−^CD117^+^ ILC progenitors in adult tonsil and lung, cord blood, and fetal liver have been reported to develop into NK cells as well as other ILCs (14, 26, 30), indicating that the ILC subsets can diverge at CD34^−^ stages. In addition, adult CD56^bright^ NK cells can express low levels of RORγt (28). We therefore analyzed whether CD34^−^ fetal cells harbored cells with both NK cells and ILC3 characteristics, and, if so, in which organs they reside. We identified Eomes^+^ NK cells expressing low levels of RORγt in fetal liver, intestine, and lung (**Fig 4A**), largely confined to CD16^−^ NK cells (**Fig. 4B-C**). In contrast, RORγt^high^ ILC3s did not express Eomes (**Fig. 1A, Fig. 4A**). Whereas all Eomes^+^ subsets expressed CD161 and PLZF (**Fig. 4D-E**), the Eomes^+^CD16^−^RORγt^+^ cells more frequently expressed CD127 and CD117 and less frequently expressed CD7 and CD45RA compared to Eomes^+^CD16^−^RORγt^−^ cells and Eomes^+^CD16^+^RORγt^−^ cells (**Fig 4F-I**). Collectively, our data suggest that a subset of CD34^−^Eomes^+^ cells within fetal tissues have a phenotype consistent with being derived from a RORγt^+^ progenitor.

**Figure 4.**
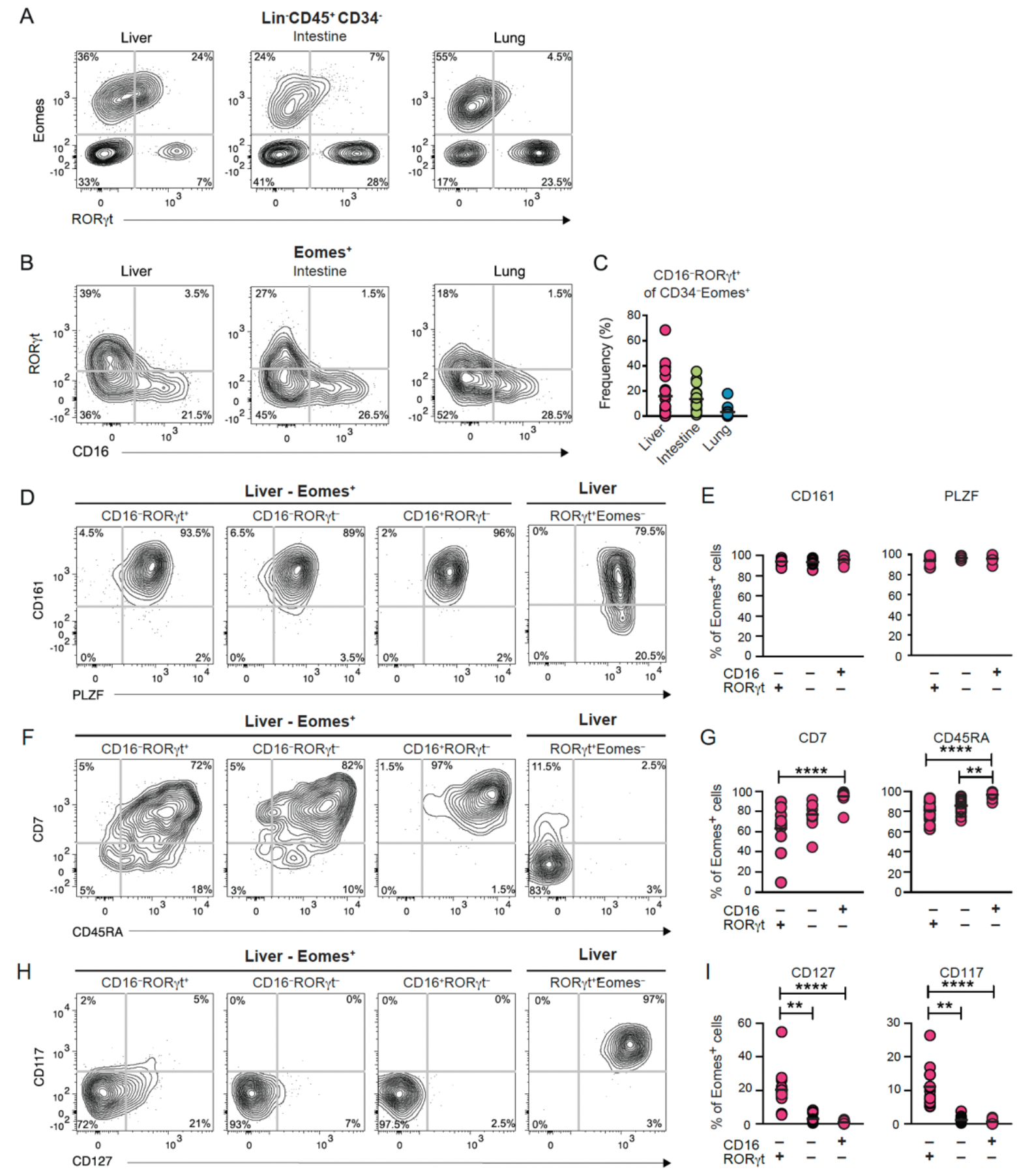
A subset of CD34^−^CD16^−^ cells co-express RORγt and Eomes. (**A**) Representative contour plots for Eomes and RORγt expression by Lin^−^CD45^+^CD34^−^ cells in matched fetal liver, intestine, and lung at PCW 8. (**B**) Representative contour plots for RORγt and CD16 expression by Lin^−^CD45^+^CD34^−^Eomes^+^ cells in liver, intestine, and lung. (**C**) Frequency of CD16^−^RORγt^+^ of CD34^−^Eomes^+^ cells in liver (*n* = 23), lung (*n* = 16), and intestine (*n* = 17). PCW 7-20. (**D-I**) Representative contour plots and summary of frequencies of (**D-E**) CD161 (*n* = 15) and PLZF (*n* = 5), (**F-G**) CD7 (*n* = 11) and CD45RA (*n* = 15), and (**H-I**) CD117 (*n* = 15) and CD127 (*n* = 15) on Eomes^+^CD16^−^RORγt^+^, Eomes^+^CD16^−^RORγt^−^, Eomes^+^CD16^+^RORγt^−^, and Eomes^−^RORγt^+^, respectively in fetal liver (PCW 7-10). Statistical comparison by non-parametric ANOVA. \**padj* < 0.05, \*\**padj* < 0.01, \*\*\**padj* < 0.001, \*\*\*\**padj* < 0.0001.

## Discussion

Here we demonstrate that mature human fetal ILCs are present in multiple tissues as early as post-conception week 6, with distinct subset composition, kinetics, and gene expression patterns between tissues. We further identify putative ILC progenitors, identified as CD34^+^RORγt^+^Eomes^+^ and CD34^+^RORγt^+^Eomes^−^ cells, exclusively in fetal intestine.

RORγt^+^ ILC3s with lymphoid tissue inducer function (LTi cells) have been shown to be crucial for the development of lymphoid structures in mice, including lymph nodes and Peyer’s patches (38). Consistent with an LTi-like function, human RORγt^+^ ILC3 can induce expression of ICAM and VCAM on mesenchymal stem cells (MSC) (6, 12). Our results show that mature ILC3s, expressing high levels of RORγt and being predominantly CD45RA^−^CD7^−^, are present across organs weeks prior to the development of lymphoid tissues. CD304^+^ ILC3s also expressed high levels of *LTB* and *LTA* in all tissues, consistent with an LTi-like function. Although the role ILC3s play in non-lymphoid tissues in the developing fetus remains unclear, their expression of *IL17A* and *IL22* suggest a potential involvement in regulating a controlled inflammatory milieu during fetal development. This may contribute to protecting against intrauterine infections and preparing epithelial barriers for interactions with commensal bacteria after birth (39, 40).

NK cells were the most frequent ILC population in fetal liver, lung, and skin, and were present at a similar frequency as RORγt^+^ ILC3s in fetal intestine (Fig. 1E). High expression levels of *GZMA* and *GZMB*, as well as CD16, already in the first trimester, indicates that fetal NK cells are indeed mature and functional, as previously reported for fetal NK cells in 2^nd^ trimester tissues (19). Early expression of CD16 on fetal NK cells suggests a potential role in providing protection against infections, possibly aided by maternally derived antibodies. NK cell-mediated protection against infections might be particularly important at the early stages of fetal immune development, where T cells are largely absent in the first trimester, and robust antibody responses by fetal B cells only emerge towards the end of the second trimester (41).

Our results revealed differences in ILC subset composition both between tissues and over gestational stages. Within each ILC subset we discovered tissue-specific gene expression patterns, including well known regulators of ILC function. For example, fetal liver CD304^+^ ILC3s expressed higher levels of *IL7R* and *PTPN6*, suggesting a less differentiated state and heightened response to IL-7 and stem cell factor (SCF). In contrast, CD304^+^ ILC3s in lung and intestine may be more differentiated and prone to establish tissue residency based on their expression of *NCR2, ALOX5AP, RGS1, RGS2*, and *ZNF331*. Similarly, CD16^+^ NK cells in fetal liver expressed higher levels of *CCNB1*, associated with cell division, and *IFNG*, while displaying lower expression levels of KIR genes and *GZMB*, suggesting a less differentiated state compared to CD16^+^ NK cells in other fetal tissues. Notably, both CD304^+^ ILC3s and NK cells in all tissues expressed high levels of genes typically linked to their respective functions, such as *LTA* and *LTB* in CD304^+^ ILC3s, and granzymes in NK cells. Combined, these findings indicate that distinct ILC subsets in different fetal tissues may respond differently to a given stimulus.

The observed tissue-specific gene expression patterns may be due to imprinting by the local tissue environment, selective recruitment of mature ILCs, or tissue-restricted ILC development. Tissue-specific gene expression patterns were almost completely unique to each ILC subset (Fig. 2B) and therefore argue against a universal tissue imprint on all immune cells in a given tissue. However, it remains possible that each ILC subset can respond differently to the same tissue environment. Furthermore, selective recruitment could not be ruled out, as there was likely heterogeneity even within the relatively homogenous subsets we studied here.

The observed tissue-specific gene expression patterns could be due to ILC development from tissue-restricted ILC progenitors giving rise to transcriptionally distinct ILC subsets between different tissues. Indeed, the identification of CD34^+^RORγt^+^Eomes^+/−^ cells in intestine, but not in other tissues, supports the notion of tissue-specific ILC progenitors. However, whether these progenitors solely give rise to ILCs in intestine or if they possess capacity to seed ILCs to other tissues, *e*.*g*. the lung, as suggested for mouse ILC2s after pathogen challenge (42), remains unknown.

Previous studies have identified CD34^+^RORγt^+^ ILC-restricted progenitors in secondary lymphoid tissues (28, 29), but not in peripheral blood, cord blood, thymus, or bone marrow (28). In line with the phenotype of CILPs (CD34^+^CD45RA^+^CD117^+^IL-1R1^+^ cells) in adult tonsil (28), most CD34^+^RORγt^+^ cells in fetal intestine expressed CD7, CD45RA, and CD117. However, there was considerable variability in expression of e.g. CD127, CD161, and CD10 in the CD34^+^RORγt^+^Eomes^−^ and CD34^+^RORγt^+^Eomes+ populations, highlighting the heterogeneity within even well-defined populations of potential ILC progenitors. Whether the distinct CD34^+^RORγt^+^Eomes^−^ and CD34^+^RORγt^+^Eomes^+^ populations represent a developmental trajectory from a multipotent ILC progenitor to a more NK cell restricted progenitor or whether they represent two distinct progenitors remains unclear.

Beyond the CD34^+^ ILC progenitors, CD34^−^ ILCPs in cord blood, adult blood, lung, tonsil, and fetal liver contain multipotent progenitors capable of differentiating into ILC1, ILC2, and ILC3s, suggesting late diversification in ILC differentiation (14, 26). RORγt expression in Eomes^+^CD94^+^CD16^−^ NK cells, along with increased CD127 and CD117 expression compared to RORγt^−^Eomes^+^CD94^+^ NK cells, suggest a gradual commitment to the NK cell lineage from a RORγt^+^ progenitor. In contrast to the previously defined CD34^−^CD7^+^CD117^+^CD127^+^ ILC progenitors in adults (14), the vast majority of ILC3s lacked CD7 and CD45RA in first trimester fetal tissues, although expression increased gradually in the second trimester (Fig. S1D). Whether the CD7^−^ ILC3s early in fetal development represent a distinct wave of ILC3 development, which is subsequently replaced by CD7^+^ ILC3s, remains unknown. This would, however, be consistent with a layered immune development as proposed for mouse and human fetal B cells and T cells (43, 44).

Our results establish a foundation for further exploration of the heterogeneity of human ILCs and their progenitors across fetal tissues, using for example *in vitro* and *in vivo* fate mapping strategies.

### Data limitations and perspective

We have identified putative CD34^+^RORγt^+^Eomes^+/−^ progenitors in the fetal intestine. We were however limited in further exploring the differentiation capacity of these cells in vitro, as fixation and permeabilization of the cells would be required to detect intracellular expression of Eomes and RORγt, precluding isolation of live cells.

In our analysis of tissue-dependent gene expression, we sorted relatively homogenous ILC subpopulations. However, there is likely heterogeneity even within these subsets, which in turn limited our ability to determine whether preferential recruitment of subpopulations or imprinting by the tissue environment underlies the observed tissue-dependent gene expression patterns we observe.

Future studies in larger number of donors using single cell RNA sequencing and single cell *in vitro* differentiation assays, could be helpful in further dissecting the tissue-dependent transcriptional patterns and ILC developmental pathways.

## Material and methods

### Human tissues and blood

First and second trimester fetal tissues were obtained from the Developmental Tissue Bank core facility at the Karolinska Institutet (PCW 6-12) and the Women’s Options Center at San Francisco General Hospital (second trimester, gestational weeks 15-22), respectively. For first trimester samples, post conceptional week was determined using ultrasound, anatomical landmarks and actual crown-rump-length.

For second trimester samples post conceptional week was estimated by measuring foot-pad size. Tonsils were collected from adult patients (20–65 years of age) undergoing tonsillectomy because of obstructive sleep apnea syndrome.

To isolate cells, tissues were cut into smaller pieces, and incubated for 30 min at 37°C with 0.25mg/ml collagenase 2 (Sigma-Aldrich), passed through a 70um nylon mesh, and diluted with R10 media (RPMI1640 with 10% FCS, penicillin, streptomycin, and L-glutamine). For first trimester samples 0.2mg/ml DNase (Roche) was also added during the digestion. Isolated cells were either stained directly or cryopreserved in FCS with 10% DMSO and stored in liquid nitrogen until use. Fetal liver and adult tonsil samples used for analysis of NKp80 expression (Fig. S1C) were not digested using collagenase to avoid any potential effects of enzymatic activity.

### Flow cytometry

Antibodies and clones against the following proteins were used (clone, fluorophore, company): CD3 (UCTH1, PECy5, Beckman Coulter), CD7 (Horizon V450 or BUV395, BD Biosciences), CD10 (HI10a, APC-Cy7, Biolegend), CD14 (M5E2, Horizon V500, BD Biosciences, or BV510, Biolegend), CD16 (3G8, APC-Cy7 or BUV496, BD Biosciences), CD19 (HIB19, Horizon V500, BD Biosciences, or BV510, Biolegend), CD34 (581, ECD, Beckman Coulter, or PE-Dazzle, Biolegend), CD45 (HI30, Alexa700, BioLegend, or BUV805, BD Biosciences), CD45RA (MEM-56, Qdot655, Invitrogen, or HI100, BV785, Biolegend), CD94 (#131412, custom-biotinylated, RnD Systems), CD117 (104D2D1, PE-Cy5.5, Beckman Coulter), CD127 (R34.34, PE-Cy7, Beckman Coulter, or A019D5, BV785, Biolegend), CD161 (HP-3G10, Brilliant Violet 605 or PerCP, BioLegend), CD304 (12C2, BV421, Biolegend), Eomes (WF1928, FITC, or eF660, eBioscience), ROR𝛾t (AFKJS-9, PE, eBioscience or Q21-559, PE or BV650, BD Biosciences), GATA-3 (TWAJ, eF660, eBioscience, or L50-829, PE-Cy7, BD Biosciences), T-bet (4B10, BV421, Biolegend, or O4-46, PE-CF-594, BD Biosciences), PLZF (#6318100, APC, RnD Systems, or Mags.21F7, PE, eBioscience), NKp80 (4A4.D10, PE or PE-Cy7, Miltenyi), CRTH2 (BM16, Horizon V450, BD Biosciences).

Secondary staining steps were performed using streptavidin Qdot585, Qdot605, Qdot 655 (Invitrogen), or streptavidin BV650 (Biolegend). All samples were stained with Live/Dead Aqua (Invitrogen). After cell surface staining with antibodies diluted in FACS wash (PBS with 2% FCS and 2mM EDTA), cells were fixed using FoxP3 Transcription Factor staining kit (Invitrogen) for 30 min and incubated with antibodies against intracellular proteins for 30 min, and resuspended in FACS wash before analyses using BD LSRII SORP, BD LSR Fortessa or BD Symphony flow cytometers. Data was analyzed using FlowJo v10 (BD Biosciences).

### Statistical analysis of flow cytometry data

For matched samples with non-gaussian distribution we used Friedman’s test with Dunn’s multiple comparisons test. For matched samples with gaussian distribution we used one-way ANOVA with Geisser-Greenhouse correction and Holm-Sidak’s multiple comparison test. Statistical analyses were performed using Prism software version 8.0 (GraphPad Software Inc).

### Cell sorting and RNA sequencing

Thawed cryopreserved cells from fetal liver, lung, intestine and skin (n = 3, PCW 9.5) were stained with the following antibodies (clone, fluorophore, company): CD3 (UCHT1, FITC, Biolegend), CD7 (M-T701, BUV395, BD Biosciences), CD14 (61D3, PE-Cy5, Invitrogen), CD15 (W6D3, PE-Cy5, Biolegend), CD16 (3G8, APC-Cy7, BD Biosciences), CD19 (HIB19, PE-Cy5, Biolegend), CD34 (581, ECD, Beckman), CD45 (HI30, AlexaFluor 700, Biolegend), CD94 (HP-3D9, BB700, BD Biosciences), CD117 (104D2, BUV737, BD Biosciences), CD127 (HIL-7R-M21, BV421, BD Biosciences), CD304 (U21-1283, BUV661, BD Biosciences), and live/dead Aqua (Invitrogen) for 30 min at 4°C, followed by washing in FACS wash. NK cells and ILC3s were first enriched by gating on Live CD45^+^CD3^−^CD14^−^CD15^−^CD19^−^CD34^−^ cells and sorting CD94^+^ and CD94^−^CD127^+^ cells into separate tubes. From the enriched cells, three subsets were sorted: CD94^+^CD16^−^ NK cells and CD94^+^CD16^+^ NK cells from the CD94^+^ enriched cells and CD94^−^CD127^+^CD304^+^ ILCs from the CD94^−^CD127^+^ enriched cells (**Fig. S2A**). For each subset from each organ 35-100 cells were sorted in duplicates into 4.2 µl of lysis buffer (0.2% Triton X-100, 2.5 µM oligo-dT (5′-AAGCAGTGGTATCAACGCAGAGTACT30VN-3′), 2.5 mM dNTP, RNAse Inhibitor (Takara)) in a 96-well V-bottom PCR plate (Thermo Fisher). Before further processing cells were stored at -80°C. RNA libraries were prepared using the standard SmartSeq2 protocol with 18 PCR cycles for cDNA amplification. The cDNA quality was assessed by bioanalyzer (Agilent, High Sensitivity DNA chip). 2 nanograms of amplified cDNA were used for our custom tagmentation protocol and indexed with Nextera XT primers. Samples were pooled and sequenced on a NextSeq2000 with P3 100 flow cell.

### Transcriptome analysis

Following sequencing, basecalling and demultiplexing was done using bsl2fastq (v2.20.0.422) with default settings generating Fastq files for further downstream mapping and analysis. Alignment was conducted using STAR 2.7.5b to GRC38/hg38.101 genome sequence from ensemble. Count reads in exons were generated with featureCounts (v1.5.1) The resulting count matrix was analyzed using DESeq2(v.1.38.3) in R(v.4.2.2). In short, raw counts from duplicates were averaged and used for input into DESEq2 and normalized. For each cell subset, only genes expressed by ≥ 2 samples with ≥50 counts were used for further analysis. Variance stabilizing transformation was used to transform the data, and limma (v.3.54.2) was used to correct for donor batch effects. Differentially expressed genes were determined by adjusted p-value < 0.01 and log2-fold change > 1 by pairwise comparison between tissues (intestine vs liver, lung vs intestine, lung vs liver, skin vs intestine, skin vs liver, and skin vs lung). Z-scores for differentially expressed genes and additional canonical markers were calculated and used for input in pheatmap (v.1.0.12) to generate heatmaps. For tissue specific differentially expressed genes, only genes uniquely differentially upregulated in one tissue for each subset were selected and visualized. Euler plots were generated using all differentially expressed genes and the nVennR(v.0.2.3) package. The adjusted p values from the differential gene expression analysis from the DESeq2 results were used for showing significance in plots for genes of interest. For each subset, genes used for visualization (heatmaps, euler plots and gene expression plots) had to be present in at least 2 donors from the same tissue with counts greater than 200.

## Ethical approval

Tissue samples were collected after informed consent and with the approval from either the Swedish Ethical Review Authority, or the University of California San Francisco (UCSF) Committee on Human Research.

## Data availability

RNA-seq count matrices and meta data with accession number xxx are available at Gene Expression Omnibus https://www.ncbi.nlm.nih.gov/geo/). (Will uploaded at time of publication). (GEO; be

## Author contribution

IER, MAI, and JaM designed the study, performed experiments, analyzed data, made figures, and wrote the manuscript. LL collected and processed fetal tissue and provided technical expertise. JEM performed experiments. DF provided adult tonsil tissue. JeM, NKB, and DFN provided input on study design. NM performed experiments, provided input on study design, and reviewed data. All authors provided input on the manuscript before submission.

## Acknowledgements

We thank the patients and staff at the Women’s Options Center at UCSF, San Francisco General Hospital and the KI Developmental Tissue Bank at Karolinska Institutet for providing prenatal tissue, and the MedH Flow core facility for providing cell sorting.

---

**Supplementary figure 1:**
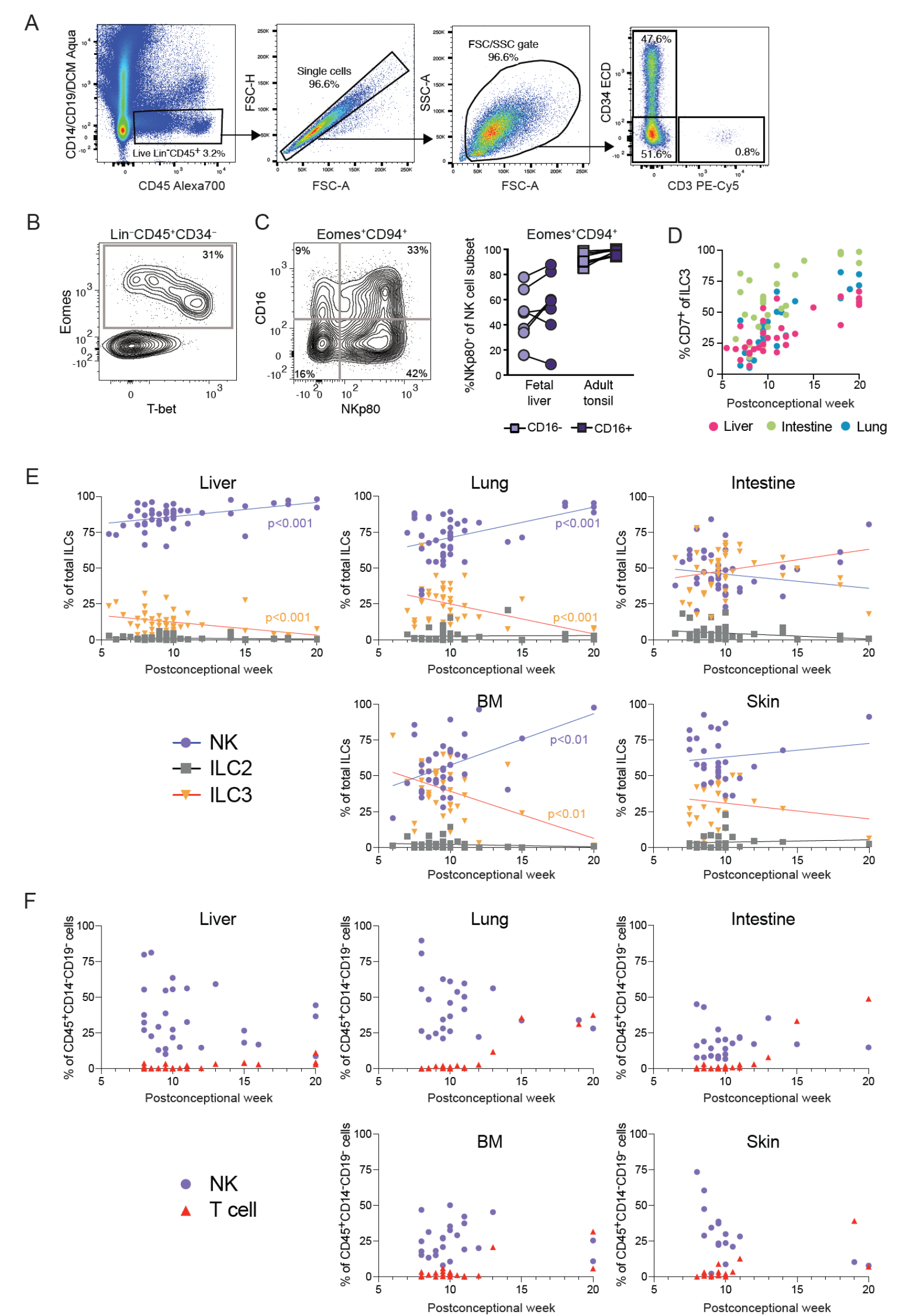
Gating strategy and lymphoid cell frequencies over gestational age. (**A**) Representative gating strategy for CD45^+^CD14^−^CD19^−^CD3^−^CD34^+^ cells, CD45^+^CD14^−^CD19^−^CD3^+^ T cells and CD45^+^CD14^−^CD19^−^CD3^−^CD34^−^ cells (PCW 8). (**B**) Representative contour plot of Eomes and Tbet expression in Lin^−^CD45^+^CD34^−^ cells in fetal liver at PCW 8. (**C**) Representative contour plot of NKp80 and CD16 expression on fetal Eomes^+^CD94^+^ NK cells in fetal liver (left) and frequency of NKp80^+^ cells of CD16^−^ and CD16^+^ NK cells in fetal liver (*n* = 7, PCW 7-14) and adult tonsil (*n* = 9) (right). (**D**) Frequency of CD7^+^ cells among ILC3s over gestational age in fetal liver (pink, n= 31), intestine (green, n=28), and lung (blue, n=27). (**E**) Frequency of NK cells (purple), ILC2s (gray) and ILC3s (orange) of total ILCs over gestational age in fetal liver (*n* = 53), lung (*n* = 47), intestine (*n* = 47), bone marrow (BM, *n* = 44) and skin (*n* = 29). (**F**) Frequency of NK cells (purple) and T cells (red) of total CD45^+^CD14^−^CD19^−^ cells across gestational age in fetal liver (*n* = 32), lung (*n* = 28), intestine (*n* = 26), BM (*n* = 27), and skin (*n* = 17).

**Supplementary figure 2.**
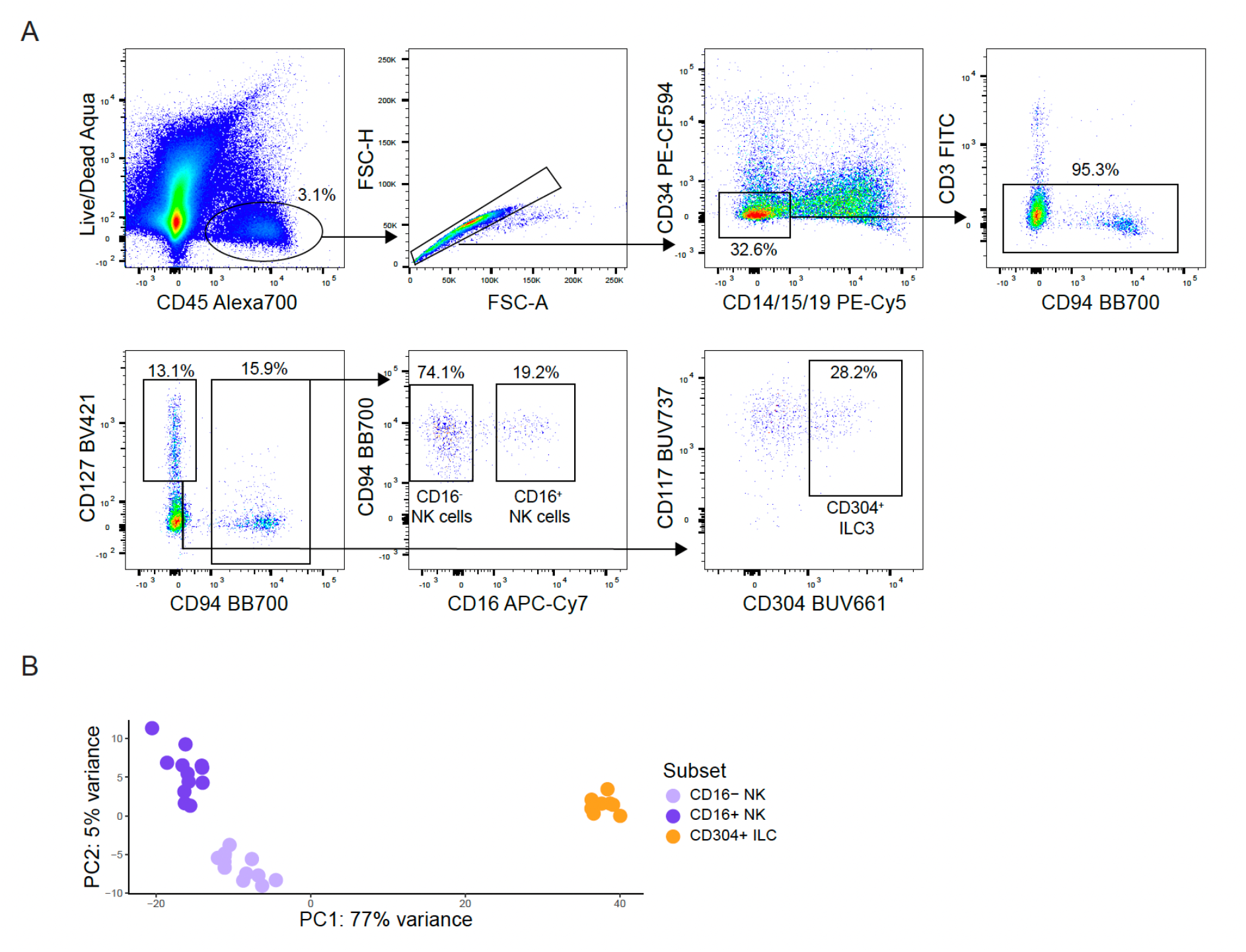
Gating strategy for sorting ILC subsets for RNA sequencing. (**A**) Gating strategy for FACS-sorting of CD45^+^CD14^−^CD15^−^CD19^−^CD3^−^ CD34^−^CD94^+^CD16^+^, CD45^+^CD14^−^CD15^−^CD19^-^ CD3^−^CD34^−^CD94^+^CD16^−^ NK cells, and CD45^+^CD14^−^CD15^−^CD19^−^CD3^−^CD34^−^CD94^−^CD127^+^CD304^+^ ILC3s. Representative plots from fetal intestine (PCW 9.5). (**B**) Principal component analysis (PCA) of donor batch corrected CD16^−^ NK cells (light purple), CD16^+^ NK cells (dark purple), and CD304^+^ ILC3s (orange) based on RNA expression. Colored by subset identity, *n* = 2-3 with cells from 4 tissues each.

**Supplementary figure 3.**
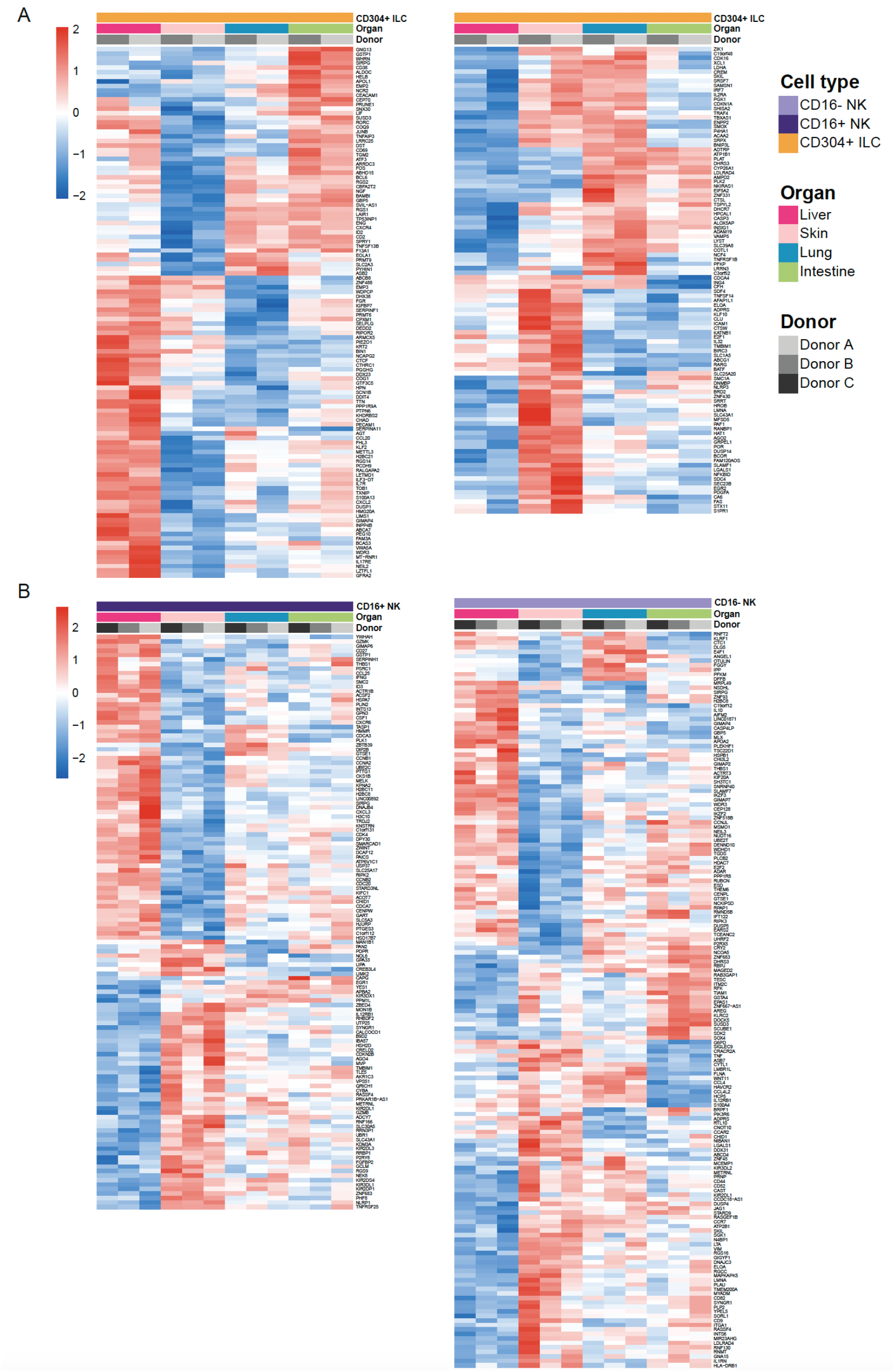
Differentially expressed genes between tissues for ILC subsets. (**A-B**) Heatmaps showing z-scores of all differentially expressed genes, clustered hierarchical by row, from pairwise comparison between fetal tissues for CD304^+^ ILC3 (orange, *n* = 2) (**A**) CD16^−^ cells (light purple, *n* = 3) and CD16^+^ NK cells (dark purple, *n* = 3) (**B**). Bars annotate cell type, organ, and donor identity (padj<0.01, log2 fold change>1).

